# Cryo-EM structure of disease-related prion fibrils provides insights into seeding barriers

**DOI:** 10.1101/2021.08.10.455830

**Authors:** Qiuye Li, Christopher P. Jaroniec, Witold K. Surewicz

## Abstract

One of the least understood aspects of prion diseases is the structure of infectious prion protein aggregates. Here we report a high-resolution cryo-EM structure of amyloid fibrils formed by human prion protein with Y145Stop mutation that is associated with a familial prion disease. This structural insight allows us not only to explain previous biochemical findings, but also provides direct support for the conformational adaptability model of prion transmissibility barriers.

---

Prion diseases are a group of transmissible neurodegenerative diseases in which the infectious agent is a misfolded aggregate of a prion protein, PrP^1–3^. Human prion diseases may be sporadic, acquired by infection, or have a hereditary origin due to mutations in the PrP-encoding gene. One of the most intriguing disorders of the latter class is a cerebrovascular amyloidosis associated with the Y145Stop mutation^4,5^. Amyloid fibrils formed in vitro by mouse Y145Stop PrP fragment (or PrP23-144) have been recently shown to be infectious, causing transmissible disease in mice^6^. Furthermore, this C-terminally truncated PrP variant has been extensively used as a model for studying molecular aspects of prion propagation, providing important insights into the mechanisms of transmissibility barriers and emergence of new prion strains^7–9^. However, interpretation of these findings at the structural level has been hampered by the lack of high-resolution information about the architecture of PrP23-144 fibrils. Here, we bridge this critical gap, by determining a cryo-EM structure of human PrP23-144 (huPrP23-144) fibrils at a near-atomic resolution.

huPrP23-144 amyloid fibrils were prepared as described in the Methods section. Inspection of cryo-EM data and subsequent 2D classification revealed only one type of fibrillar aggregates in our sample. The ordered amyloid core of these fibrils has a diameter of ∼10 nm, with the remaining, largely disordered parts of huPrP23-144 wrapping around the amyloid core and contributing to a larger apparent diameter of ∼20 nm (Extended Data Fig. 1a and b). Atomic force microscopy imaging revealed that fibrils are characterized by left-handed twist (Extended Data Fig. 1c).

Helical reconstruction of the cryo-EM data allowed us to determine a density map for the ordered amyloid core with a nominal resolution of 2.86 Å (Fig. 1a and b, Extended Data Fig. 2 and Extended Data Table 1). This core shows a parallel, in-register architecture, in which each fibril consists of four identical protofilaments with a C2 symmetry (Fig. 1b and c). Four-protofilament fibrils are rather unusual, even though they have been reported for a fragment of TDP-43 low complexity domain^10^ and recombinant SAA protein^11^. A near-atomic model was built for the fibril core that maps to residues 108-141 (Fig. 1d and e). The identity of this core region is similar to that suggested by previously reported solid-state NMR studies, though the fibril samples in those studies appear to be composed of two protofilaments^9,12^. In our cryo-EM-based model, each subunit has an “S” shape, largely extended backbone geometry and encompasses three relatively short β-strands (residues 109-112, 133-135, and 138-140) (Fig. 1c and e). The 113-125 region is rich in rigid turns, with numerous solvent-inaccessible hydrophobic residues participating in intramolecular interactions involving side chains of M112/A116/A117 and A115/A118/V121/L125 (Fig. 1d and f), in line with those observed by solid-state NMR^9,12^. Another cluster of solvent-inaccessible hydrophobic side chains maps to the C-terminal part of the core and includes M129, A133, I139, and F141 (Fig. 1d and f).

**Fig. 1:**
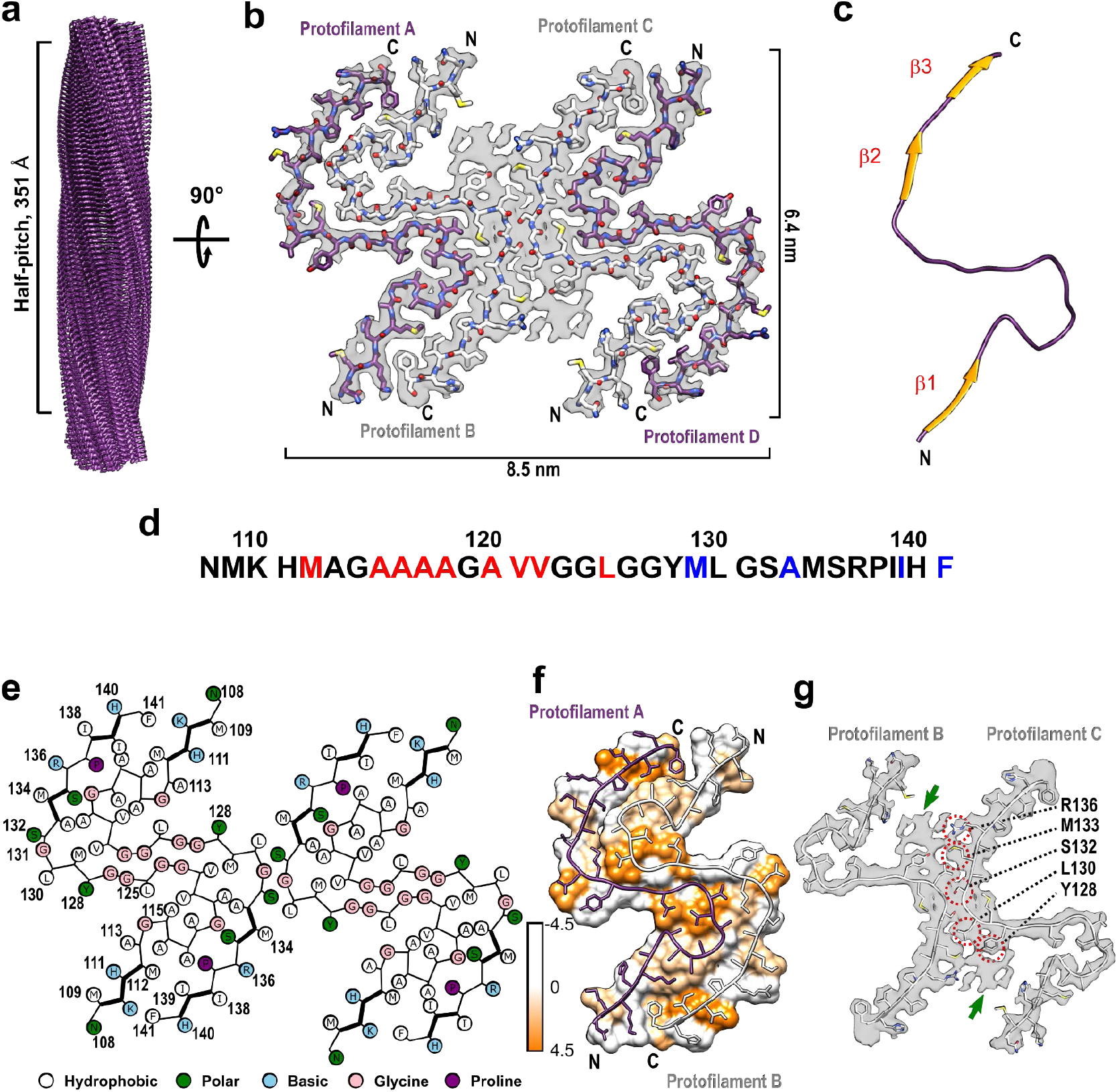
Cryo-EM structure of huPrP23-144 amyloid fibrils. **a** Cryo-EM density map showing a left-handed twisted helix with a half-pitch of 351 Å and a classical parallel, in-register β-sheet architecture. **b** The top view of the atomic model superimposed on the cryo-EM density map. The amyloid core contains four identical protofilaments. Two outer protofilaments with greater accessibility to water are depicted in purple; two inner protofilaments are depicted in white. **c** One representative subunit of the fibril core with β-strands shown in orange. **d** Amino acid sequence for the highly ordered core of huPrP23-144 fibrils. N- and C-terminal solvent-inaccessible hydrophobic residues are shown in red and blue, respectively. **e** Schematic representation of one cross-sectional layer of the amyloid core, with β-strands shown as thicker lines. **f** Hydrophobicity of the cross-section of protofilaments A and B, with hydrophobicity levels colored according to Kyte-Doolittle^19^ (top view). **g** One cross-sectional layer of protofilaments B and C shown as atomic model superimposed on the cryo-EM map (top view). Extra unassigned densities (likely representing side chains of residues outside the core region) are indicated by green arrows; residues within the amyloid core involved in the inter-protofilament interactions are indicated by red circles.

These two clusters of hydrophobic resides play a key role in stabilizing the four-protofilament structure of the fibrils. In this structure, protofilaments A and B (as well as C and D) are arranged in a head-to-tail manner, with a large dry interface between them that is stabilized largely by intermolecular interaction between the N-terminal hydrophobic residues (M112, A116, A117, A120, and V122) and C-terminal hydrophobic residues (M129, A133, I139, and F141) (Fig. 1e and f). Interestingly, no classical steric zipper motif is observed within this dry interface. The above-described intermolecular interactions, together with intramolecular interactions involving hydrophobic residues within the 112-125 region, result in a structure in which most (13 out of 18) hydrophobic side chains are buried within the amyloid core interior. The interface between inner protofilaments B and C is much shorter, with intermolecular hydrogen bonds involving only L130 and S132. The latter interface is additionally stabilized by side chain-side chain interactions involving nearby residues Y128, L130, M133, and R136 (Fig. 1e and g).

Protofilaments A and B (and C and D) assembled through large hydrophobic interfaces are arranged in such a manner that N- and C-terminal hydrophobic residues within each monomeric subunit interact with two different subunits in the adjacent protofilament, as illustrated in Extended Data Fig. 3. Such a non-planar assembly of subunits results in rugged surfaces of fibril ends, with solvent exposure of several hydrophobic side chains. Specifically, at the top end of the fibril, two clusters of N-terminal hydrophobic residues (M112/A116/A117 and A120/V122) are exposed to the solvent. Depending on the stage of fibril elongation, these residues are either from protofilaments B and C (Fig. 2a) or protofilaments A and D (Fig. 2b). Similarly, two clusters of hydrophobic C-terminal residues (M129/A133 and I139/F141) from protofilaments A and D (Fig. 2c) or B and C (Fig. 2d) are exposed at the bottom end of the fibril. These exposed hydrophobic residues likely play a key role in the recruitment and conformational conversion of the incoming monomers.

**Fig. 2.**
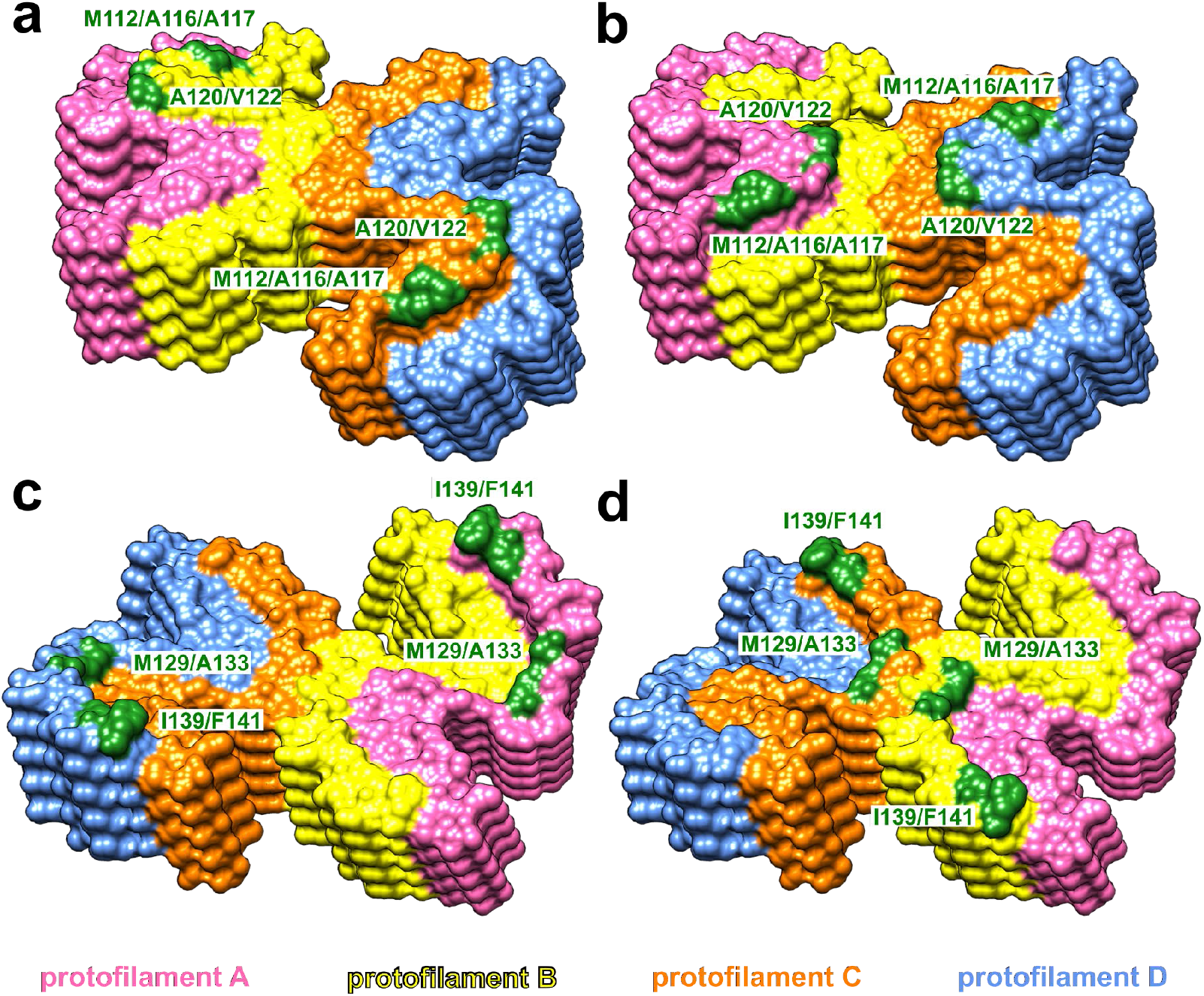
Solvent-exposed amino acid residues at the top and bottom ends of huPrP23-144 fibrils. At the top end of the fibril, two clusters of hydrophobic residues (M112/A116/A117 and A120/V122) in protofilaments B and C (yellow and orange, **a**) or protofilaments A and D (pink and blue, **b**) are exposed to the solvent. At the bottom end of the fibril, two clusters of hydrophobic residues (M129/A133 and I139/F141) in protofilaments A and D (pink and blue, **c**) or protofilaments B and C (yellow and orange, **d**) are exposed to the solvent.

The present structural data have a number of important implications for understanding the propagation and seeding specificity of huPrP23-144 amyloid fibrils. First, earlier studies revealed that neither huPrP23-137 nor huPrP23-139 fragments are able to form amyloid fibrils (even in the presence of PrP23-144 amyloid seeds), and that the shortest C-truncated variant of huPrP23-144 showing fibril forming capacity is huPrP23-141^13^. This may be rationalized by the present high-resolution structural model, as this model reveals that the C-terminal hydrophobic residues I139 and F141 are directly involved in the dry interface between the protofilaments, playing an important role in fibril-stabilizing interactions.

Second, one of the most puzzling aspects of prion diseases is the phenomenon of species-dependent transmissibility barriers. Animal experiments^14^ and studies in vitro using the PrP23-144 model^7,8^ have led to the hypothesis that the key determinant of prion transmission is the ability of monomeric PrP substrate of the host to adapt to the structure of the donor prion aggregate. However, in the absence of high-resolution structural data, this model remains largely conceptual in nature. As part of our studies using the PrP23-144 system, we have established that there is a strong cross-seeding barrier between huPrP23-144 and Syrian hamster PrP23-144 (ShaPrP23-144), and that this barrier is fully controlled by the identity of just one amino acid residue at position 139 (Ile and Met in huPrP and ShaPrP, respectively)^8,9^. The present data allows us to rationalize this puzzling observation in structural terms, since the replacement of Ile139 with a longer Met side chain would likely disallow ShaPrP23-144 to adapt to the structure of the huPrP23-144 seed due to steric clashes between side chains of M139 and M112 in the adjacent subunits (Extended Data Fig. 4 and 5). Thus, this structural insight provides direct, molecular-level support for the “conformational adaptability” model of prion transmissibility barriers.

Finally, it should be noted that recent studies reported the structures of (non-infectious) amyloid fibrils formed by the recombinant full-length huPrP^15,16^ as well as brain-derived infectious hamster prions^17^. For obvious reasons, the present structure is fundamentally different, as the disease-related huPrP23-144 lacks the entire C-terminal part of the prion protein. More intriguingly, the structure of huPrP23-144 fibrils is also very different from that of fibrils formed by a non-physiological huPrP90-178 fragment^18^, even though the core regions of both fibrils are similar. This strongly suggests that residues outside the core region play an important role in conformational conversion of the prion protein, and thus, caution should be exercised when interpreting structural data for fibrils formed by short, non-physiological fragments of PrP.

## Methods

### Protein expression, purification, and fibril formation

huPrP23-144 was expressed and purified as described previously^13,20^. Lyophilized protein was re-suspended at a concentration of 400 μM in 50 mM potassium phosphate buffer containing 0.01% NaN_3_, pH 6.5. Initially, we attempted to prepare fibrils in an identical way as for previous solid-state NMR studies^9,12^, by incubating the solution of monomeric protein at 25°C under quiescent conditions for 7 days. However, cryo-EM images of the end product showed heavily clumped fibrils that were not suitable for high-resolution structural studies. Therefore, to prepare a better dispersed sample, these fibrils were sonicated and then used to seed a diluted solution (40 μM) of monomeric huPrP23-144, again under quiescent conditions. The final product of the seeded reaction was briefly sonicated and, when examined by cryo-EM, showed relatively well-dispersed fibrils that were immediately used for cryo-EM data collection.

### Atomic force microscopy

For AFM analysis, 20 μl of fibril suspension was deposited on freshly cleaved mica substrate and incubated for 5 min. The surface was then washed three times with Milli-Q H_2_O and dried under N_2_. The images were obtained by NanoScope 9.1 using scan assist mode and a silicon probe (spring constant, 40 newtons/m) on a Bruker multimode atomic force microscope equipped with Nanoscope V controller.

### Cryo-EM

Three hundred mesh lacey carbon grids (Ted Pella) were coated with 0.1 mg/ml graphene oxide followed by 0.1% poly-lysine as described previously^21,22^. Three microliters of freshly sonicated huPrP23-144 fibril suspension (40 μM) was applied to the coated grid, blotted for 7 s, and plunge-frozen in liquid ethane using a Vitrobot Mark IV (ThermoFisher Scientific). Movies were collected on a Titan Krios G3i microscope (ThermoFisher Scientific) equipped with a BioQuantum K3 camera (Gatan, Inc.), with 0.414 Å/pixel in super resolution mode (53 e^−^/Å^2^ total dose, and 48 total frames). A total of 3,604 movies were automatically collected using SerialEM^23^ with 7 shots per position. Beam image shift was applied, and defocus range was between −0.8 and −1.5 μm.

### Data processing

Movies were first corrected for drifting and binned by a factor of 2 using MotionCor2^24^. Contrast Transfer Functions (CTF) were estimated by CTFFIND 4.1.3^25^. All further processing was carried out using RELION 3.1^26–28^. Individual fibrils were manually picked, and 1,089,468 overlapping segments were extracted using an inter-box distance of 10 Å and a box size of 512 pixels. Segments were first subjected to three rounds of reference-free two-dimensional (2D) classification using T = 8 and K = 200 to remove poorly defined classes, resulting in 349,164 segments contributing to clear 2D averages. These segments were then used for subsequent three-dimensional (3D) classification employing an initial model of a featureless cylinder generated by relion_helix_toolbox^27^. The initial helical rise (4.70 Å) was calculated from the 2D class layer line profile, and the initial helical twist (−2.4°) was calculated from the crossover distance. Two rounds of three-dimensional (3D) classification were performed using K = 8 and T = 4, resulting in 46,803 segments that contributed to a high-resolution reconstruction. The reconstructed map showed a C2 symmetry, which was imposed in the subsequent processing. After 3D classification, segments were re-extracted from micrographs with a box size of 256 pixels and then subjected to iterative high-resolution gold-standard 3D refinement, Bayesian polishing^28^, and CTF refinement^29^ to reconstruct the final high-resolution map. The overall resolution was calculated to be 2.86 Å from Fourier shell correlations at 0.143 between two independently refined half-maps. Refined helical symmetry (twist = −2.41°, rise = 4.71 Å) was imposed on the post-processed map for further model building.

### Model building

An initial model was built in Coot^30^ using Glycine-rich region of residues 121-130 as a guide. Five cross-sectional layers (20 chains) were built at the central region of the density map. The model was then subjected to iterative real-space refinements in PHENIX^31,32^. β-strands were identified manually from the density map and such secondary structure restraints were implemented in the real-space refinements. After real-space refinement, side-chain orientations were further adjusted in Coot^30^ to ensure energy-favored geometry. The final model was validated using the comprehensive validation method in PHENIX^33^. A map containing twenty subunits was extracted manually from the reconstructed map using the UCSF Chimera package^34^ to calculate Fourier shell correlations between the map and the atomic model in PHENIX^33^.

### Reporting summary

Further information on experimental design is available in the Nature research Reporting Summary linked to this article.

### Data Availability

Cryo-EM density map and the atomic model of human PrP23-144 fibrils has been deposited to the Electron Microscopy Data Bank and Protein data bank with the accession codes EMD-24514 and 7RL4, respectively. Other data are available from the corresponding author upon reasonable request.

## Acknowledgements

This work was supported by NIH grants GM094357 and NS103848. We thank Kunpeng Li for help with acquisition of cryo-EM data. We are grateful to the Cryo-EM Core at CWRU School of Medicine (especially Dr. Sudha Chakrapani and Dr. Kunpeng Li) for the access to cryo-EM instrumentation.

## Competing interests

The authors declare no competing interests.

## Extended Data

**Extended Data Fig. 1.**
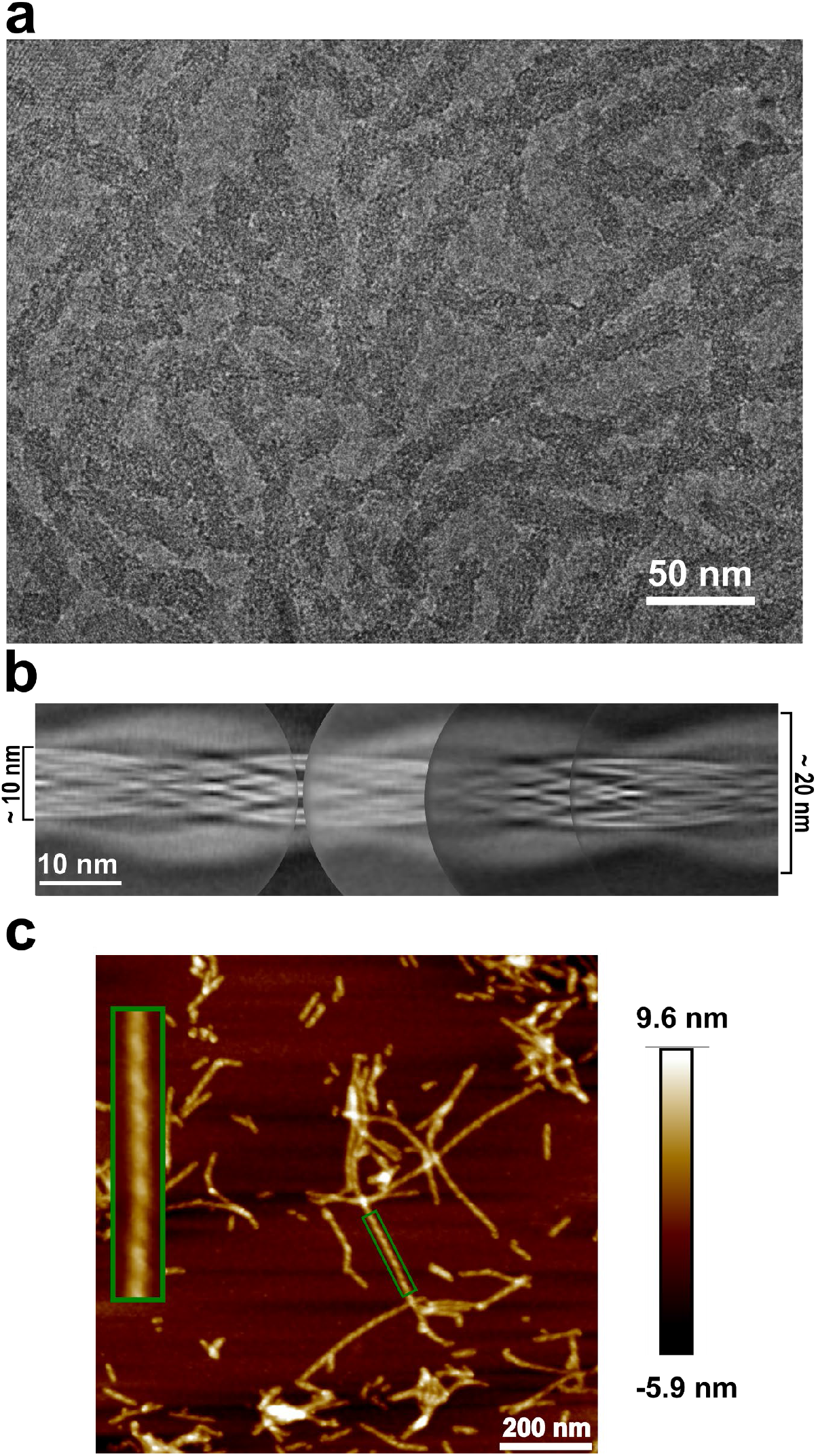
Morphology of huPrP23-144 amyloid fibrils. **a** Cryo-EM image showing one single type of morphology with an apparent twist. **b** Manually assembled full pitch of huPrP23-144 fibrils from multiple 2D class averages. The entire fibril has a thickness of ∼20 nm and the highly ordered amyloid core has a thickness of ∼10 nm. **c** AFM image of huPrP23-144 fibrils. One representative fibril in the green box is enlarged to show the left-handed twist.

**Extended Data Fig. 2.**
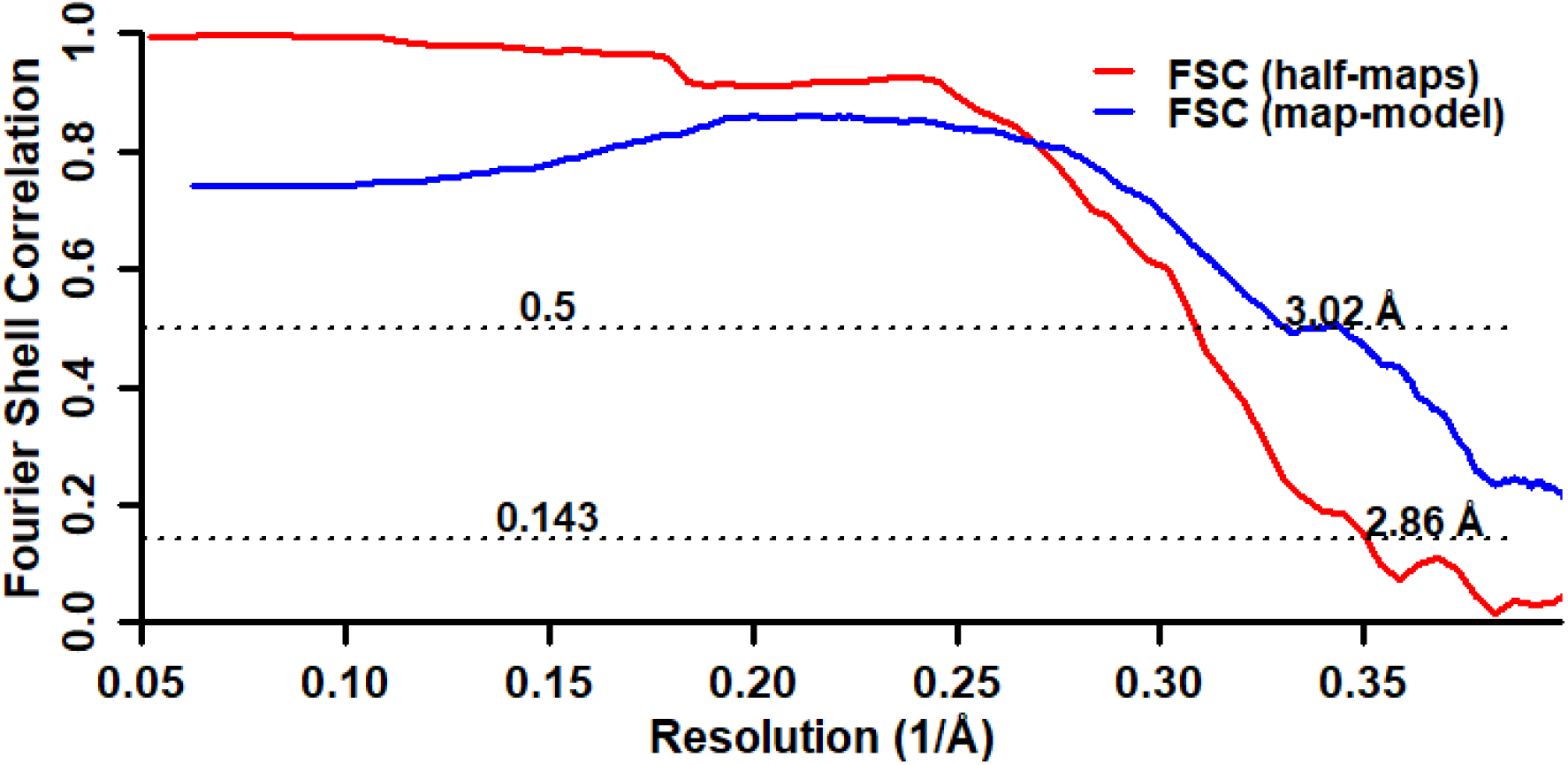
Fourier shell correlation curves between two independently refined half-maps (red) and between the map and the atomic model (blue).

**Extended Data Fig. 3.**
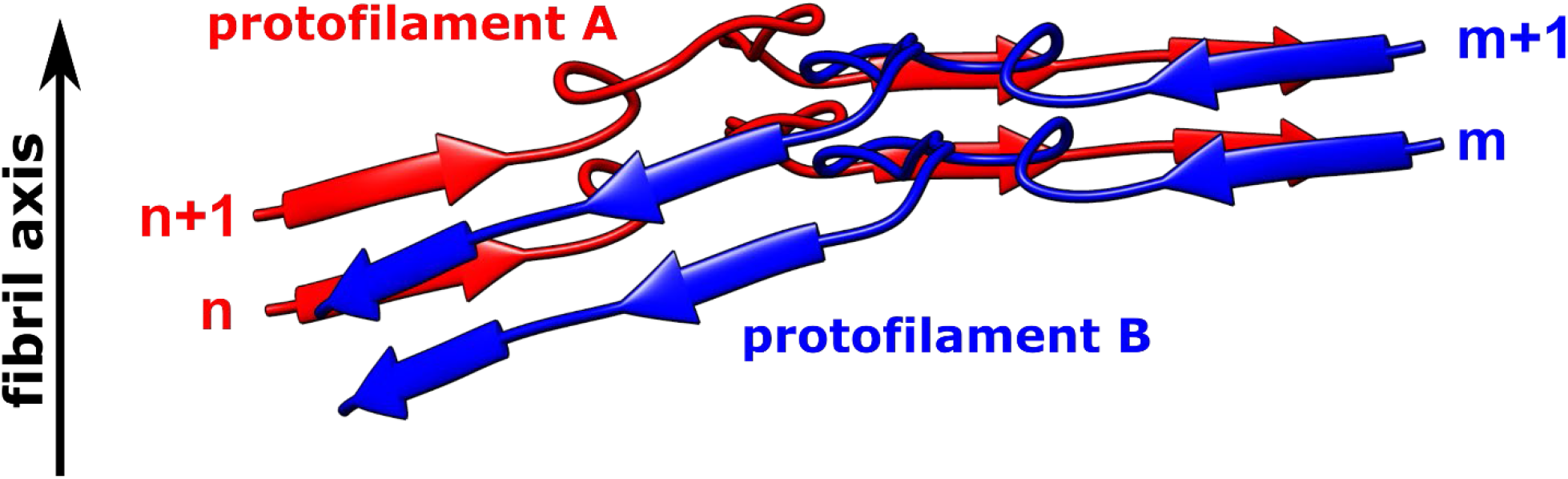
Non-planar architecture of two representative protofilaments assembled through large hydrophobic interfaces. Subunit n in protofilament A interacts with two subunits in protofilament B (subunits m and m+1); subunit m+1 in protofilament B interacts with two subunits in protofilament A (subunits n and n+1). Such a non-planar conformation results in rugged surfaces at fibril ends, with N-terminal residues at the top end and C-terminal residues at the bottom end exposed to water. A similar non-planar assembly is observed for subunits in protofilaments C and D.

**Extended Data Fig. 4.**
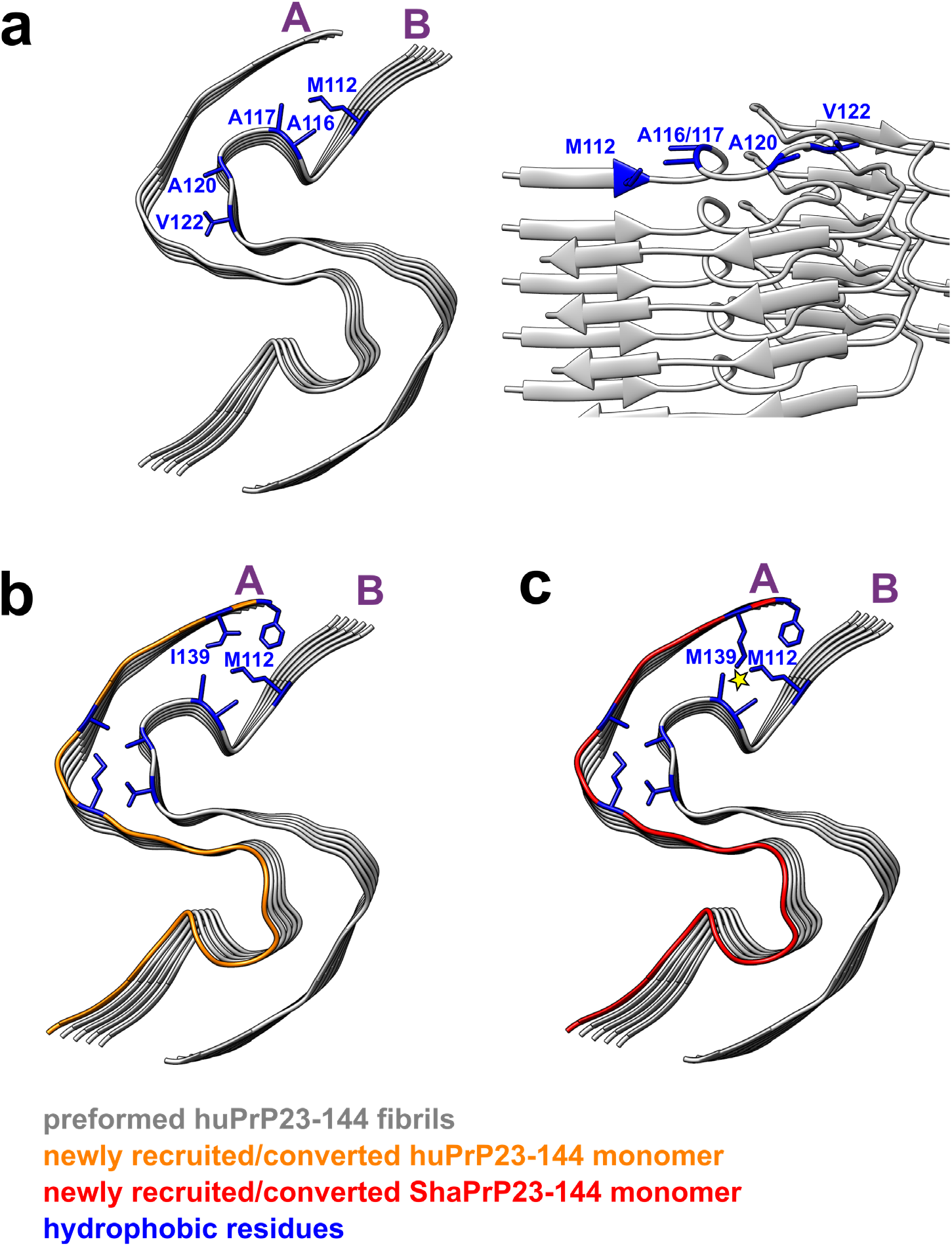
The structural model illustrating seeding reaction at the top end of huPrP23-144 fibrils in the presence of huPrP23-144 and ShaPrP23-144 substrates. Two representative protofilaments (A and B) are shown only. **a** Top and side views of a preformed huPrP23-144 fibril (seed) with solvent-exposed hydrophobic side chains shown in blue. **b** Top view of a preformed huPrP23-144 fibril (grey) with a newly recruited and converted subunit of huPrP23-144 (orange). **c** Top view of a preformed huPrP23-144 fibril (grey) with a newly recruited subunit of ShaPrP23-144 (red). Adaptation of ShaPrP23-144 to the structure of huPrP23-144 seed would lead to significant intermolecular steric clashes between bulky, elongated side chains of M112 and M139 (as indicated by the yellow star), explaining a cross-seeding barrier.

**Extended Data Fig. 5.**
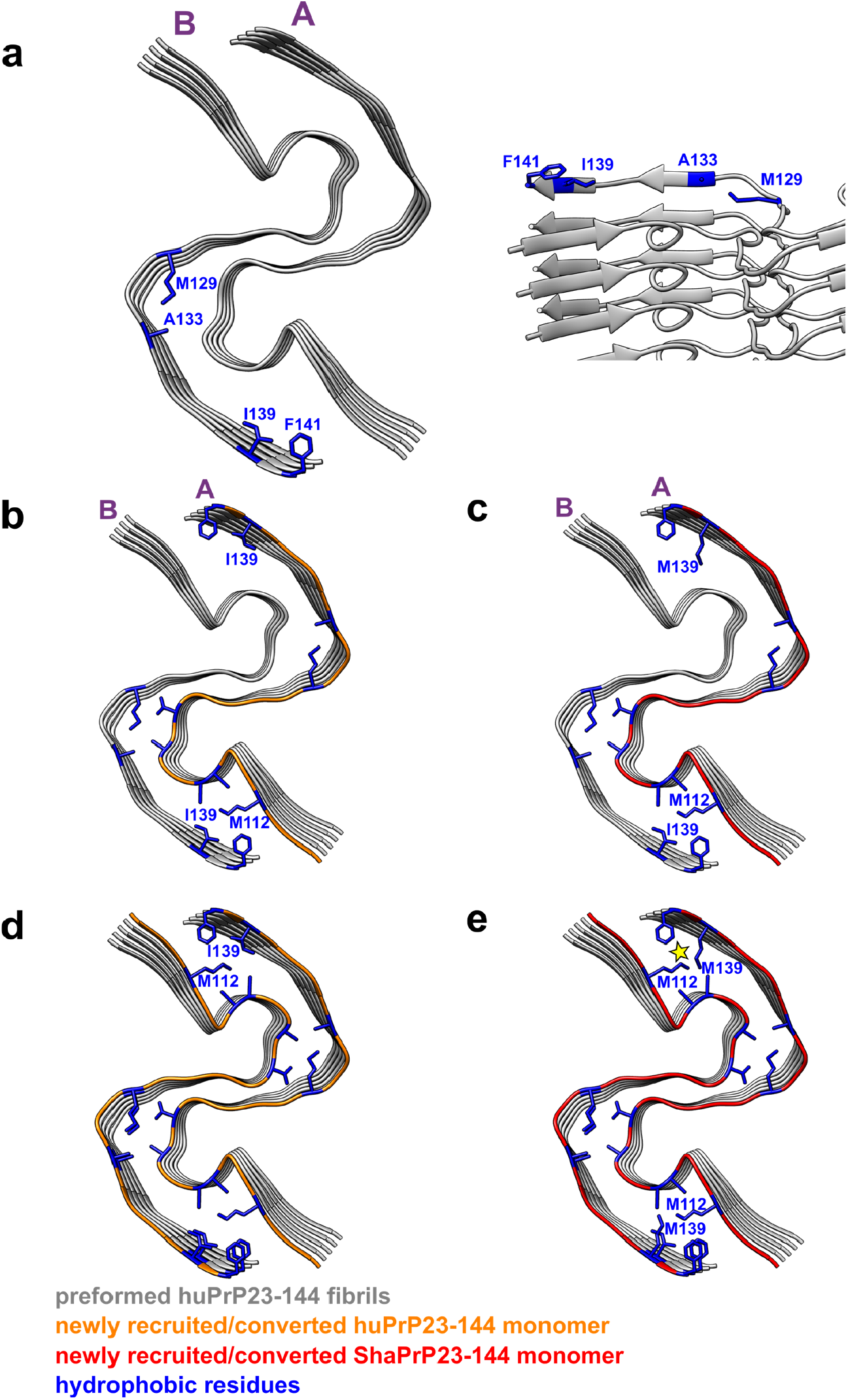
The structural model illustrating seeding reaction at the bottom end of huPrP23-144 fibrils in the presence of huPrP23-144 and ShaPrP23-144 substrates. Two representative protofilaments (A and B) are shown for illustrative purposes. **a** Bottom and side views of a preformed huPrP23-144 fibril (seed) with solvent-exposed hydrophobic side chains shown in blue. **b** Bottom view of a preformed huPrP23-144 fibril (grey) with a newly recruited (to protofilament A) and converted subunit of huPrP23-144 (orange). **c** Bottom view of a preformed huPrP23-144 fibril (grey) with a newly recruited (to protofilament A) and converted first subunit of ShaPrP23-144 (red). Due to non-planar structure, C-terminal hydrophobic residues at this end of protofilament B are protruding to water. Thus, recruitment of the first ShaPrP subunit would not result in any steric clashes. **d** Bottom view of a preformed huPrP23-144 fibril (grey) with a second newly recruited (to protofilament B) and converted subunit of huPrP23-144 (orange). **e** Bottom view of a preformed huPrP23-144 fibril (grey) with a second recruited (to protofilament B) subunit of ShaPrP23-144 (red). Adaptation of this subunit to the structure of the huPrP23-144 seed would lead to intermolecular steric clashes between side chains of M139 and M112 (as indicated by the yellow star), explaining a cross-seeding barrier.

**Extended Data Table 1.** Cryo-EM data collection, refinement, and validation statistics.

|  |  |  |
| --- | --- | --- |
| **Data collection and processing** |  |  |
| Magnification |  | × 105,000 |
| Voltage (kV) |  | 300 |
| Electron dose (e<sup>-</sup>/Å<sup>2</sup>) |  | 53 |
| Defocus range (μm) |  | -0.8 to -1.5 |
| Pixel size (Å) |  | 0.828 |
| Symmetry imposed |  | C2 |
| Initial particles images (no.) |  | 1,089,468 |
| Final particle images (no.) |  | 46,803 |
| Map Resolution (Å) |  | 2.86 |
| FSC threshold |  | 0.143 |
| **Refinement** |  |  |
| Initial model used |  | De novo |
| Map sharpening B factor (Å<sup>2</sup>) |  | -79.39 |
| Model composition |  |  |
| Non-hydrogen atoms |  | 4,620 |
| Protein residues |  | 680 |
| B factors (Å<sup>2</sup>) |  |  |
| protein |  | 88.57 |
| R.M.S.D |  |  |
| Bond lengths (Å) |  | 0.015 |
| Bond angles (°) |  | 1.390 |
| MolProbity score |  | 2.20 |
| Clashscore |  | 11.94 |
| Poor rotamers (%) |  | 0 |
| Ramachandran plot |  |  |
| Favored (%) |  | 87.50 |
| Allowed (%) |  | 12.50 |
| Disallowed (%) |  | 0 |

## Notes

### Competing Interest Statement

The authors have declared no competing interest.

